# Phosphoserine aminotransferase SerC is a central metabolic checkpoint and druggable vulnerability in *Mycobacterium tuberculosis*

**DOI:** 10.64898/2026.03.27.714911

**Authors:** Michael J Perret, Tom A Mendum, Daniel Kim, Jade Seng, Brian Robertson, Rebecca Winsbury, Simon Clark, Johnjoe McFadden, Khushboo Borah Slater

## Abstract

Serine metabolism is fundamental to the pathogenicity of *Mycobacterium tuberculosis* (Mtb), the causative agent of tuberculosis (TB), yet its full metabolic scope and therapeutic potential remain unclear. Here we show that the phosphoserine aminotransferase *serC* is an essential metabolic node that coordinates central carbon and nitrogen flux and enables intracellular survival in the host. Deletion of *serC* caused severe growth defects across macrophage and murine infection models and rewired central carbon metabolism, reducing glycolytic, tricarboxylic-acid-cycle, and methylcitrate-cycle fluxes while altering one-carbon, branched-chain, and amino-acid biosynthetic pathways. Transposon sequencing identified *sdaA*-dependent serine deamination and the glycine cleavage system as key determinants of serine-based nitrogen assimilation, while revealing redundancy in serine transport. Our findings validate serine biosynthesis as a vulnerable, druggable metabolic target in Mtb and highlight it as a promising route for the development of urgently needed therapies against TB, which continues to kill millions of people each year.

## Introduction

Tuberculosis (TB), caused by *Mycobacterium tuberculosis* (Mtb), remains one of the world’s deadliest infectious diseases, responsible for more than 1.5 million deaths annually^1^. The growing prevalence of drug-resistant Mtb strains has rendered current treatment regimens increasingly ineffective, posing a major threat to global health and placing substantial economic and clinical burdens on healthcare systems^2^. There is an urgent need for new therapeutic strategies, including novel antimicrobials active against drug-resistant Mtb and host-directed therapies (HDTs) capable of enhancing immune-mediated clearance of infection^3^. The COVID-19 pandemic further exacerbated the strain on TB diagnosis, treatment, and management, adding complexity to an already challenging global health problem. The current landscape of drug resistance and associated morbidities emphasizes the urgent need to identify new vulnerable targets and accelerate the development of next-generation TB therapies with novel mechanism of action.

Mtb is an intracellular pathogen that replicates within macrophages in a phagosome which is characterized by limited nutrient availability. To survive in this nutrient-restricted environment, Mtb must acquire and assimilate multiple host-derived nutrients. During intracellular growth, Mtb utilizes both carbon and nitrogen sources provided by the host cell^4^. Nitrogen is a fundamental building block of cellular biomass and is essential for the intracellular growth of Mtb. Previous work demonstrated that Mtb uses the asparagine transporter *ansP2* and the secreted asparaginase *ansA* to hydrolyze asparagine, release ammonia, and resist phagosomal acidification^5^. Deletion of *ansA* attenuates Mtb *in vivo*, highlighting the link between nitrogen metabolism and virulence.

Mtb preferentially utilizes amino acids such as glutamate, aspartate, asparagine, and glutamine over inorganic nitrogen sources^6^. Our previous work using stable isotope tracing and ^15^N-flux spectral analysis revealed that intracellular Mtb uses multiple host amino acids including glutamate, glutamine, aspartate, alanine, glycine, valine, and leucine within human macrophages^4^. Dual ^13^C-^15^N-metabolic flux analysis established glutamate as the central node of nitrogen metabolism and highlighted flexible nitrogen utilization strategies by Mtb to support its survival within the human host^4,7^. Together, these studies demonstrate that Mtb possesses a versatile and adaptive nitrogen metabolic network that supports intracellular survival and virulence.

Despite this progress, our understanding of nitrogen metabolism in Mtb remains incomplete. The identities and functions of many transaminases and amino acid transporters essential for intracellular survival are poorly defined. The roles of several nitrogen-related pathways remain uncharacterized, and the extent to which specific nitrogen sources influence Mtb’s physiology, virulence, and drug tolerance is not fully understood. In our previous work, we showed that serine (Ser) is not available to intracellular Mtb and that *de novo* serine biosynthesis is essential for Mtb’s growth and virulence^4^. Deletion of phosphoserine transaminase *serC* (Rv0884c), a key enzyme in Ser biosynthesis, results in Ser auxotrophy. This Δ*serC* mutant is significantly attenuated in macrophages, revealing Rv0884c as a druggable vulnerability within Mtb’s nitrogen metabolic network that can be exploited for therapeutic development.

Ser metabolism plays a fundamental role across diverse biological systems, influencing microbial physiology, antimicrobial susceptibility, and cancer cell proliferation. In bacteria, Ser-derived compounds and serine-dependent enzymes have emerged as promising therapeutic targets. Ser-based amphiphilic surfactants exhibit potent activity against antibiotic-resistant *Escherichia coli* and *Staphylococcus aureus* by disrupting bacterial membranes while remaining non-cytotoxic to human cells^8^. Targeting Ser-related metabolic pathways has also been shown to improve sensitivity to antimicrobials. Inhibition of serine O-acetyltransferase (CysE) sensitizes Gram-negative bacteria to carbapenems^9^, and serine hydroxymethyltransferase (SHMT) inhibition suppresses *Enterococcus faecium* growth and enhances nucleoside analogue activity^10^. In mammalian systems, Ser-glycine one-carbon metabolism supports nucleotide synthesis, redox balance, and tumour progression, highlighting enzymes of this pathway such as 3-phosphoglycerate dehydrogenase and phosphoserine aminotransferase as attractive anticancer targets^11–14^.

Ser metabolism is essential for growth, survival and virulence of Mtb. Mtb synthesizes Ser through several metabolic routes, with the phosphorylated pathway involving 3-phosphoglycerate as the major source^15^. This pathway consists of three enzymes, each demonstrated to be essential for Mtb’s growth^4,16^. The first enzyme, phosphoglycerate dehydrogenase (*serA1*, Rv2996c), catalyzes the initial step of serine biosynthesis and has been identified as an essential gene in Mtb^16^. The second enzyme, phosphoserine aminotransferase (*serC*, Rv0884c), converts phosphohydroxypyruvate to phosphoserine and plays a particularly critical role during intracellular infection^4^. The final enzyme, phosphoserine phosphatase (*serB2*, Rv3042c), completes the pathway and is also essential for optimal bacterial growth ^16^. Recent studies have further emphasized the importance of Ser metabolism in Mtb’s adaptation to drug tolerance and to resist hostile host environments. Previously we used genome scale metabolic modelling coupled with differential producibility analysis to translate RNA seq datasets into metabolite level signals and identified drug associated metabolic response profiles. Ser biosynthesis emerged as a key component of nitrogen metabolism and was implicated in drug tolerance, with Ser levels elevated in bedaquiline (BDQ) and isoniazid (INH) treated cells^17^. In addition to its metabolic roles, Ser biosynthesis also contributes to immune evasion. Rv0884c was upregulated under hypoxia to produce D-serine, which suppresses IFN-γ production by CD8⁺ T cells^18^. This immune suppression reduces macrophage antimicrobial activity and supports Mtb persistence in the hypoxic intracellular environment.

Together, these studies highlight that Mtb uses Ser metabolic adaptation for intracellular survival, suppression of host immunity, and tolerance to antibiotic stress. The essentiality of *serA1*, *serC*, and *serB2* highlights the indispensable nature of Ser biosynthesis in Mtb and positions these enzymes as promising targets for the development of new anti TB therapies. Despite the essentiality of Ser metabolic genes, selective inhibition strategies against these targets remain underdeveloped, and the metabolic and physiological roles of serine metabolism are not yet fully understood.

In this study, we validated the essentiality of *serC* across multiple infection models and established its metabolic functions in Mtb using systems based multi-omic approaches. We identified additional genes required for Ser-dependent nitrogen metabolism in Mtb, thereby expanding the repertoire of potential therapeutic targets.

## Results

### Rv0884c is required for serine and glycine biosynthesis in Mtb and intracellular survival

Rv0884c encodes the phosphoserine aminotransferase *serC* in Mtb, which catalyses the transamination of 3-phosphonooxypyruvate to phosphoserine (Fig. 1A). We previously established that serine is not accessible to intracellular Mtb within human macrophages and that loss of *serC* results in Ser auxotrophy and impaired intracellular growth^4^. In our previous work, we constructed an Rv0884c deletion mutant (Δ*serC*) in the H37Rv background. Here we evaluated the fitness of the mutant across multiple infection models. Consistent with our earlier observations in THP-1 macrophages (Fig. S1A), Δ*serC* Mtb exhibited a significant intracellular growth defect in primary human macrophages and in RAW 264.7 cells (Fig. 1B–C). Infection with Δ*serC* did not compromise macrophage viability, indicating that the observed attenuation reflects bacterial metabolic insufficiency rather than host cell toxicity.

**Fig 1.**
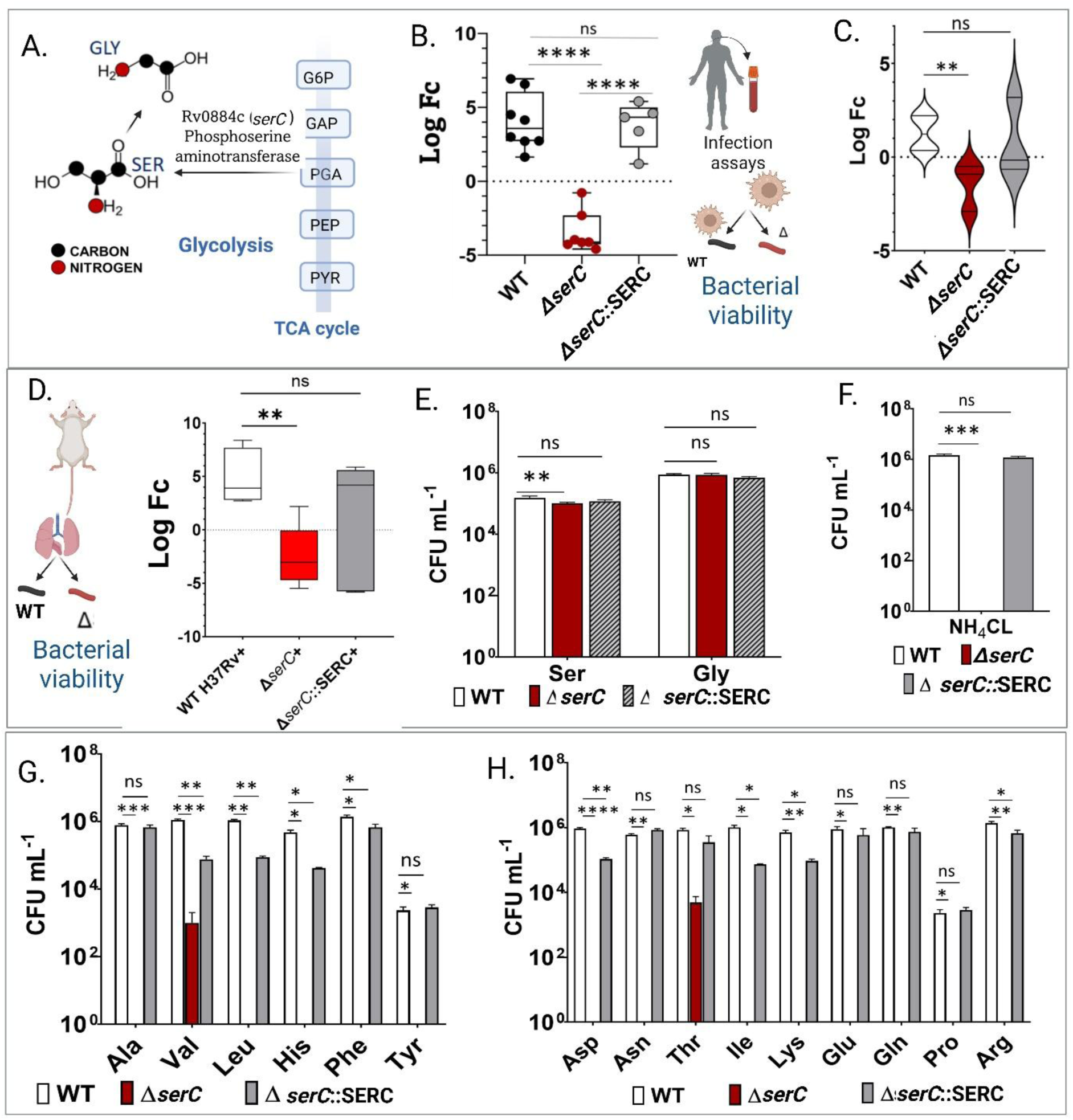
Intracellular growth and nitrogen utilization of Mtb serine auxotroph. **A)** Schematic illustrating the enzymatic role of *serC* (Rv0884c) in serine (Ser) and glycine (Gly) biosynthesis from glycolytically derived phosphoglyceric acid (PGA). **B)** Intracellular growth of wild-type (WT), Δ*serC*, and complemented Δ*serC*::serC Mtb H37Rv strains in primary human macrophages derived from PBMCs. Bacterial viability is shown as log 2-fold change (F_C_) in CFU mL^-1^ at day 7 relative to day 0. Data represent five to eight independent infections using macrophages from two donors. **C)** Intracellular growth of WT, Δ*serC*, and Δ*serC*::serC strains in murine RAW 264.7 macrophages. Log 2-F_C_ values were calculated at day 3 relative to day 0. Data represent three independent infections. **D)** *In vivo* fitness of Δ*serC* compared with WT and complemented strain in BALB/c mice. Animals were infected with 200 CFU of each strain, and lung bacterial burden was quantified at Day 1, Day 7, Day 21 and Day 28 post-infection. Five mice were used per group. Log 2-Fc values were calculated at Day 21 and Day 28 relative to Day 1. **E–H**) Nitrogen source utilization by the Δ*serC* mutant in minimal medium supplemented with individual nitrogen sources (10 mM). Growth rescue was assessed in the presence of **E**, Ser and Gly; **F**, ammonium chloride (NH₄Cl); **G**, alanine (Ala), valine (Val), leucine (Leu), histidine (His), phenylalanine (Phe), and tyrosine (Tyr); and **H**, aspartic acid (Asp), asparagine (Asn), threonine (Thr), isoleucine (Ile), lysine (Lys), glutamic acid (Glu), glutamine (Gln), proline (Pro), and arginine (Arg). Data represent the mean of three independent replicates ± SEM. * indicates statistically significant differences determined using unpaired t-test with Welch’s correction (P ≤ 0.05) or Man -Whitney test (P ≤ 0.05) for mice datasets.

To assess the *in vivo* relevance of Ser biosynthesis, we infected BALB/c mice with Δ*serC*, wild-type (WT) or complemented (Δ*serC*::SERC) Mtb. Infected animals remained clinically healthy with stable body weight throughout the experiment (Fig. S2A). Δ*serC* displayed severely reduced bacterial burden relative to the WT and complemented strain (Fig. 1D). This significant attenuation demonstrates the essentiality of *de novo* Ser biosynthesis for Mtb’s survival in the host environment. Together, these data demonstrate that Rv0884c-dependent Ser metabolism is indispensable for intracellular replication and virulence of Mtb, and highlights Ser biosynthesis as a critical metabolic vulnerability across diverse host contexts.

We next assessed the growth phenotype of the Δ*serC* mutant on a range of organic and inorganic nitrogen sources, including amino acids and ammonium chloride (NH₄Cl), to determine which substrates could rescue its Ser auxotrophy. This analysis was designed to test whether Rv0884c functions specifically in Ser metabolism. The complemented strain (Δ*serC*::SERC) served as a control to confirm that the observed phenotype resulted from loss of *serC*. Tryptophan (Trp), methionine (Met), and cysteine (Cys) did not support the growth of the WT, mutant, or complemented strains and were therefore excluded from further analysis. Among the organic nitrogen sources tested, Ser and glycine (Gly) provided the most robust rescue, restoring 66–98% of wild-type growth in vitro (Fig. 1E). In contrast, NH₄Cl, the standard inorganic nitrogen source for Mtb failed to rescue the mutant, validating the essentiality of *serC* and the non-redundant nature of the RV0884c-mediated transaminase reaction, which reinforces their potential as drug targets. Threonine (Thr) and valine (Val) supported only minimal recovery (0.09–0.6% of wild-type levels; Fig. 1G–H). As Thr is derived primarily from tricarboxylic acid (TCA) cycle intermediates and Val from pyruvate, their limited rescue suggests probable indirect metabolic connectivity with Ser biosynthesis. Together, these findings support the specificity of Rv0884c for *de novo* Ser production and reinforces its central role in nitrogen metabolism of Mtb.

### Deletion of *serC* impairs carbon and nitrogen utilisation in Mtb

To determine how the loss of *serC* affects central metabolism in Mtb, we performed *in vitro* isotopic-labelling experiments. Exponentially growing cultures were incubated with either [U-¹³C₃]-glycerol or [¹⁵N₁]-ammonium chloride, and incorporation of ¹³C and ¹⁵N into the proteinogenic amino acids were quantified by gas chromatography-mass spectrometry (GC-MS) (Fig. 2A). These experiments were designed to compare carbon (C) and nitrogen (N) assimilation between the WT and the Δ*serC* strain. Overall, total ¹³C and ¹⁵N enrichment across amino acids was significantly reduced in Δ*serC*, indicating impaired incorporation of both elements despite supplementation of serine to rescue auxotrophy (Fig. 2B). As expected, Ser and Gly showed no detectable ¹³C or ¹⁵N labelling (below natural isotopic abundance) confirming complete loss of *de novo* Ser and Gly biosynthesis in the mutant.

**Fig. 2.**
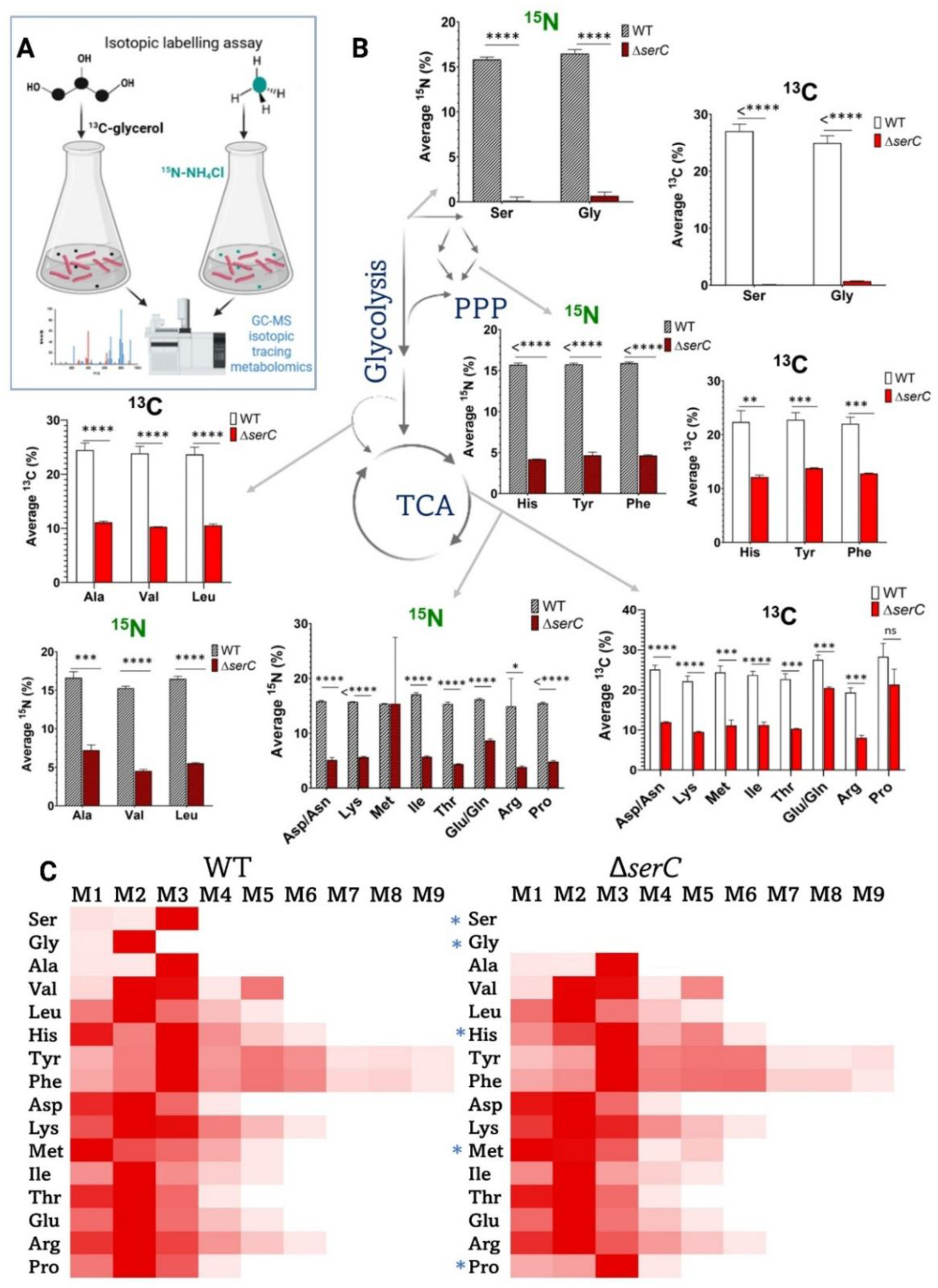
Isotopic labelling analysis reveals altered carbon and nitrogen metabolism in Δ*serC* Mtb. **A)** Schematic overview of the ¹³C and ¹⁵N isotopic-labelling workflow. WT and Δ*serC* Mtb strains were cultured in media containing either U-[¹³C₃]-glycerol or [¹⁵N₁]-ammonium chloride as single-isotope tracers. Cells were harvested at mid-exponential phase, and incorporation of ¹³C and ¹⁵N into the proteinogenic amino acids was quantified by GC–MS. All experiments were performed in biological triplicate for both strains. **B)** Mean ¹³C (%) and ¹⁵N (%) label incorporation into amino acids derived from (i) glycolysis: serine (Ser), glycine (Gly), valine (Val), alanine (Ala), leucine (Leu); (ii) the pentose phosphate pathway (PPP): histidine (His), tyrosine (Tyr), phenylalanine (Phe); and (iii) the tricarboxylic acid (TCA) cycle: aspartate (Asp), asparagine (Asn), lysine (Lys), methionine (Met), isoleucine (Ile), threonine (Thr), glutamate (Glu), glutamine (Gln), arginine (Arg) and proline (Pro). **C)** ¹³C mass isotopomer distribution (MID) profiles for amino acids in WT and Δ*serC* strains. Values represent means from three biological replicates and were used to generate the heat maps. M1–M9 denote ¹³C-labelled mass isotopomers. Data underlying all panels are provided in Data file S2.

Amino acids derived from pyruvate: alanine (Ala), valine (Val) and leucine (Leu) and from the pentose phosphate pathway: histidine (His) exhibited significantly reduced labelling in Δ*serC*, accounting approximately half of the WT levels (Fig. 2A). Among the TCA cycle-derived amino acids, all except methionine (Met) and proline (Pro) showed decreased incorporation of both isotopes. For Pro, the total ¹⁵N enrichment was reduced to less than half of the WT, whereas Met displayed unchanged ¹⁵N incorporation but significantly reduced ¹³C labelling (Fig. 2B).

To further examine carbon-backbone synthesis, we analysed ¹³C mass isotopomer distributions (MIDs) (Fig. 2C). ¹⁵N MIDs were not evaluated because single-nitrogen amino acids (for example, Ala, Ser, Gly) do not provide sufficient isotopic distribution resolution to identify pathway usage. Consistent with total labelling, ¹³C isotopomer abundances were substantially lower in Δ*serC*, reflecting a global reduction in carbon metabolic flux. Except for Ser, Gly, Met, Pro and His, MID patterns were similar between the WT and mutant. The absence of Ser and Gly isotopomers in Δ*serC* reflects the loss of their biosynthetic pathways, whereas altered MID profiles for Met, Pro and His suggest rerouting of carbon flux into these amino acids in the mutant. Together, these findings demonstrate that deletion of Rv0884c diminishes overall rates of carbon and nitrogen metabolism in Mtb and perturbs the synthesis of Ser, Gly, Met, Pro and His.

### Deletion of *serC* reprogrammes central metabolic fluxes in Mtb

To define how the loss of *serC* reshapes central carbon metabolism (CCM) in Mtb, we quantified *in vivo* metabolic fluxes in the WT and Δ*serC* strains using ^13^C-metabolic flux analysis (MFA). Our aim was to identify the metabolic nodes and fluxes perturbed by *serC* deletion and thereby establish its metabolic function. Mass isotopomer distributions (MIDs) of proteinogenic amino acids obtained from [U-^13^C_3_]-glycerol labelling were integrated into a previously established metabolic model describing carbon atom transitions in CCM^4^. Nitrogen fluxes could not be resolved due to insufficient MID information from [U-^15^N_1_]-NH_4_Cl labelling, as most amino acids contain only a single nitrogen atom and therefore lack informative isotopomer transitions for MFA. Cultures were grown in batch mode and harvested in late exponential phase, corresponding to pseudo–steady state conditions^19,20^. Carbon fluxes were normalised to a glycerol uptake rate of 100 (Fig. 3).

**Figure 3.**
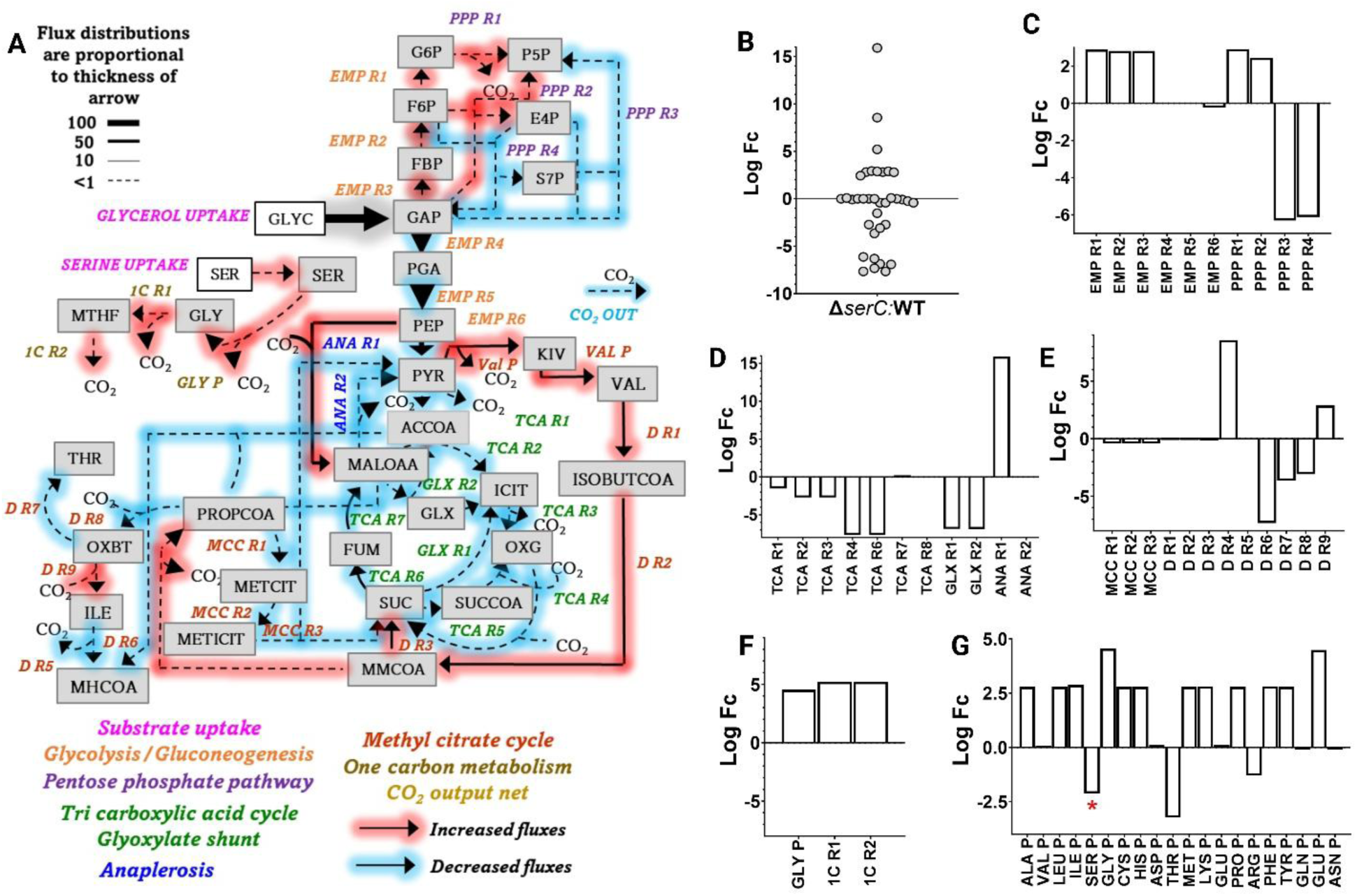
^13^C-metabolic flux analysis reveals extensive metabolic rewiring in *serC* mutant. **A)** Flux map illustrating changes in carbon flux distributions in Δ*serC* compared with WT. Fluxes were estimated using the Mtb carbon atom–transition network (Data file S3) and normalised to a total glycerol uptake rate arbitrarily set to 100. [U-^13^C_3_]-glycerol served as the labelled substrate, and ^13^C mass isotopomer distributions of protein-derived amino acids were used for flux estimation. Arrows represent fluxes through central carbon metabolism, with line width proportional to flux magnitude. Metabolite lists for flux maps are provided in Data file S3. Labelling experiments were performed in triplicate for both WT and Δ*serC*. **B)** Log 10-fold changes of all quantified fluxes in Δ*serC* relative to WT. **C)** Log 10-fold changes in glycolytic and gluconeogenic fluxes. **D)** Log 10-fold changes in TCA cycle and glyoxylate shunt fluxes. **E)** Log 10-fold changes in methylcitrate cycle (MCC) and valine degradation pathway fluxes. **F)** Log 10-fold changes in amino-acid biosynthetic fluxes. * Indicates a zero flux for *de novo* synthesis for Ser biosynthesis (SER P) in Δ*serC*. Data are mean ± standard error of the mean from three independent labelling experiments.

A substantial fraction of CCM fluxes was significantly reduced in the mutant (blue arrows, Fig. 3A, B). Glycerol enters CCM via glycolysis, and glycolytic fluxes from glyceraldehyde-3-phosphate (GAP) to pyruvate (PYR) (EMP R3-R6) were significantly lower in Δ*serC* relative to WT (Fig. 3A, C). This reduction in glycerol catabolism is a direct consequence of *serC* deletion. In contrast, gluconeogenic fluxes (EMP R1-R3) were elevated in the mutant. Because these reactions consume ATP when operating in the glycolytic direction, their elevation in the mutant may reflect an energy-conserving strategy in Δ*serC*.

Fluxes through the oxidative pentose phosphate pathway (PPP R1 and PPP R2) were also higher in the mutant (Fig. 3C). PPP R2 generates pentose-5-phosphates (ribose- and ribulose-5-phosphates), precursors for nucleotide and histidine biosynthesis. The elevated oxidative PPP flux is consistent with the altered ^13^C MID patterns observed for His, indicating PPP rewiring upon *serC* deletion. Because the oxidative PPP is a major source of NADPH for lipid biosynthesis and other anabolic processes^21^, these data suggest that Δ*serC* may experience an increased demand for reductant. In contrast, non-oxidative PPP fluxes were reduced relative to WT (Fig. 3C).

Fluxes through the TCA cycle and glyoxylate shunt were generally lower in the mutant (Fig. 3D). However, anaplerotic flux from phosphoenolpyruvate to oxaloacetate (ANA R1) was elevated, indicating enhanced CO_2_ fixation to replenish TCA intermediates. Val biosynthesis and degradation fluxes were also increased (Fig. 3E, G). Val degradation generates precursors for propionyl-CoA (PROPCOA), which is normally detoxified via the methylcitrate cycle (MCC). MCC fluxes (MCC R1-R3) were reduced in Δ*serC* (Fig. 3E), suggesting that PROPCOA is diverted toward cell-wall lipid synthesis such as phthiocerol dimycocerosates (PDIMs) and sulfolipids rather than being catabolised through the MCC^21,22^.

Fluxes towards Ser-derived metabolites, including Gly (Gly P) and one-carbon folate cycle reactions (1C R1, 1C R2), were elevated in the mutant (Fig. 3F). One-carbon metabolism supports the synthesis of nucleic acids, amino acids, creatine, and phospholipids. Deletion of serC resulted in an increased production of branched chain amino acid fluxes (VAL P, LEU P, Ile P), histidine (HIS P), asparagine (ASN P), glutamate (GLU P), proline (PRO P), methionine (MET P), tyrosine (TYR P), phenylalanine (PHE P), lysine (LYS P), alanine (ALA P) while threonine (THR P) and arginine (ARG P) fluxes were reduced.

Together, ^13^C-MFA revealed extensive metabolic rewiring in Δ*serC*, characterised by reduced glycolytic and TCA cycle fluxes, increased Val degradation, diminished MCC activity, enhanced oxidative PPP flux, and selective upregulation of amino-acid biosynthetic pathways.

### Transposon sequencing identifies genes essential for Mtb growth on serine

To identify genetic determinants required for growth of Mtb on serine, we performed transposon (Tn) library selections using minimal medium containing glycerol as the sole carbon source and either NH₄Cl or SER as the sole nitrogen source (Fig. 4A). Both selected libraries exhibited characteristics consistent with previously published Tn-seq datasets. The NH₄-selected library contained 533 genes classified as essential (ES) and 137 as growth-disadvantaged (GD), 92% of which had been previously designated ES or GD on 7H10 medium (^23^). Among genes classified as non-essential (NE) or growth-advantaged (GA), 97% were concordant with prior designations on 7H10 medium. The SER-selected library contained 504 ES and 180 GD genes (Data file S4). Several genes involved in SER metabolism including *serA1*, which catalyses the first committed step in Ser biosynthesis^24^, and *glyA1*, which converts SER to GLY were absent from both selected libraries, consistent with their previously reported essentiality for central nitrogen metabolism (Fig. 4B, C).

**Figure 4.**
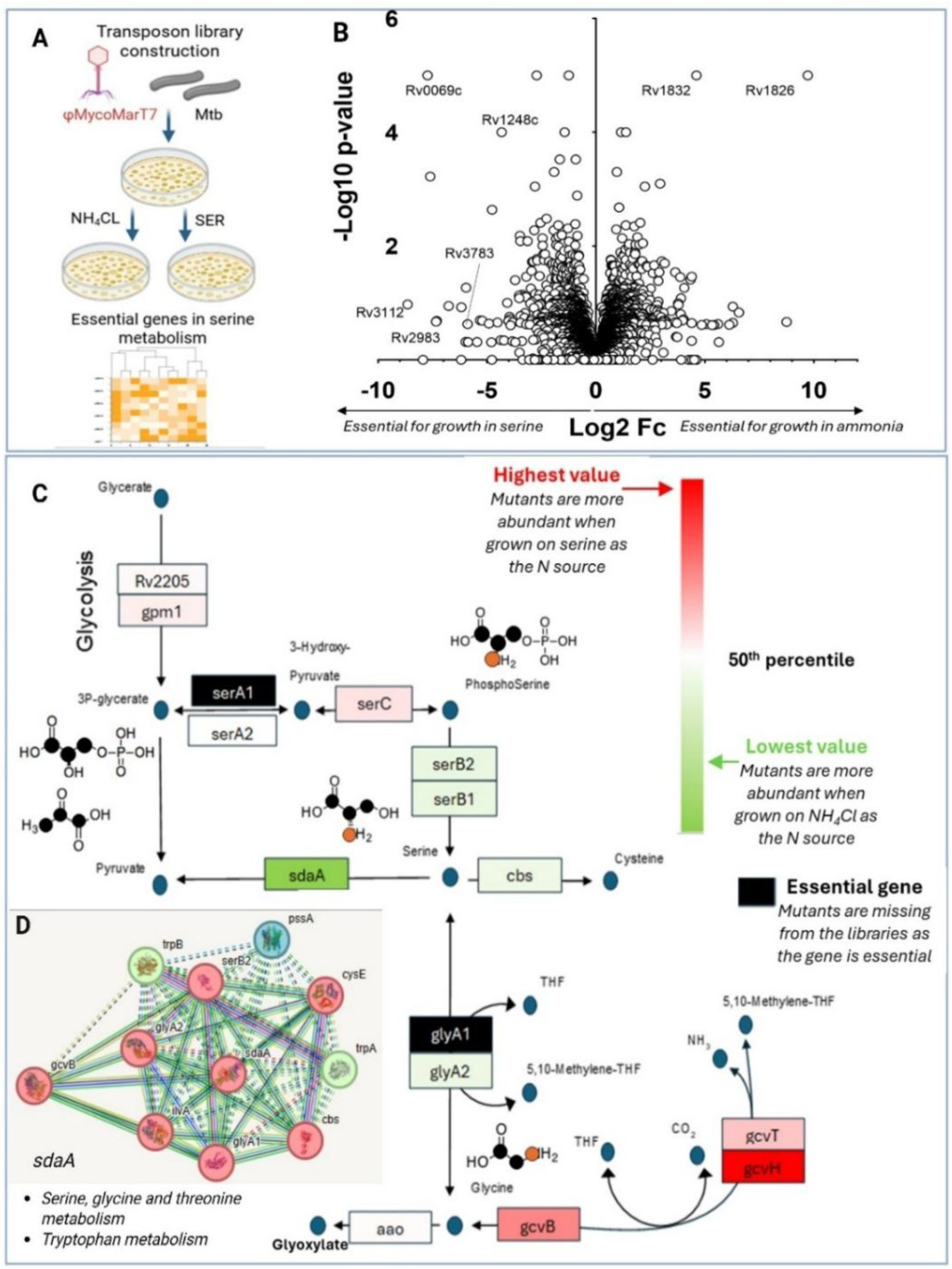
Transposon (Tn) library screening identifies genes essential for Ser metabolism in Mtb. **A)** Schematic of Tn library construction using the temperature-labile phage φMycoMarT7. Libraries were selected on minimal medium containing glycerol as the carbon source and either ammonium chloride (NH₄Cl) or serine (SER) as the sole nitrogen source. **B)** Volcano plot comparing mutant abundance in SER- versus NH₄-selected libraries. The y-axis shows the –log₁₀ *P* value, and the x-axis shows the log₂ fold change in abundance (SER vs. NH₄). **C)** Serine metabolic pathway map showing mutant abundance and gene essentiality across the two Tn libraries. Mutants enriched during growth on SER relative to NH₄ are shown in a red gradient, whereas mutants enriched on NH₄ relative to SER are shown in a green gradient. Increasing colour intensity corresponds to greater relative abundance. *serA1* and *glyA1* (black) were essential under both nitrogen conditions. **D)** STRING v12 protein–protein association network for *SdaA* (Rv0069c), illustrating functional links between genes involved in Ser, Gly, Thr and Trp metabolism. Line thickness reflects confidence in the predicted associations.

Only eight mutants exhibited significantly different fitness between the NH₄- and SER-selected libraries. The most pronounced difference was observed for *sdaA*, which encodes the primary L-serine deaminase responsible for SER catabolism (Fig. 4C). SdaA deaminates SER to generate PYR and NH₄⁺, the latter serving as a nitrogen source for biosynthesis of nucleotides and amino acids. Although threonine deaminase (*ilvA*) can deaminate Ser in some organisms^25^, its absence from the SER-selected library likely reflects its essential role in ILE biosynthesis^26^. The strong dependence on *sdaA* for growth on SER suggests that *ilvA* contributes minimally to SER deamination in Mtb, despite evidence from *M. smegmatis* that *ilvA* can use SER as a substrate *in vitro*. Because SER was the sole nitrogen source in the SER-selected medium, the loss of the primary serine deaminase would be expected to impose a substantial fitness cost, validating *sdaA* as an essential gene for serine-dependent nitrogen metabolism.

A second notable group of genes with altered fitness comprised of genes involved in the glycine cleavage system (GCS) *gcvB*, *gcvH*, and to a lesser extent *gcvT* (Fig. 4C). These genes were previously classified as essential on 7H10 medium^23^. In our dataset, GCS mutants were present and relatively enriched in the SER-selected library but not in the NH₄-selected library. The GCS catalyses glycine degradation to generate 5,10-methylenetetrahydrofolate (THF) and NADH, which feed into one-carbon metabolism and ultimately S-adenosylmethionine (SAM) synthesis. Impaired SAM regeneration is thought to be bactericidal^27^. When SER is abundant, *glyA1*-mediated conversion of SER to GLY likely produces sufficient THF to compensate for the loss of GCS activity. These findings highlight the importance of the GCS under standard growth conditions and *in vivo*, where SER availability is limited^4^.

Finally, network analysis of genes depleted in the SER-selected library revealed that proteins involved in Ser, Gly, Thr, and Trp metabolism including *serB2*, *cysE*, *glyA2*, *gcvB*, *cbs*, *ilvA*, *glyA1*, *trpA*, and *trpB* clustered with *sdaA* (Fig. 4D), highlighting a tight link between Ser biosynthesis and amino acid metabolic pathways.

### Transposon screening to identify potential serine transporters

We anticipated that the Tn-seq screen would identify candidate serine transporters, as loss of such transporters should impair the growth of Mtb on serine as the sole nitrogen source but not when NH_4_^+^ is provided. However, the only putative amino acid transporter showing a significant fitness difference between the two selective libraries was *ansP2* (Rv0346c), a known asparagine transporter (Fig. 5A). Mutants of *ansP2* were present in both libraries and were, in fact, more abundant in the SER-selected library, indicating an increased fitness on Ser which contrasts with what would be expected for a Ser transporter.

**Figure 5.**
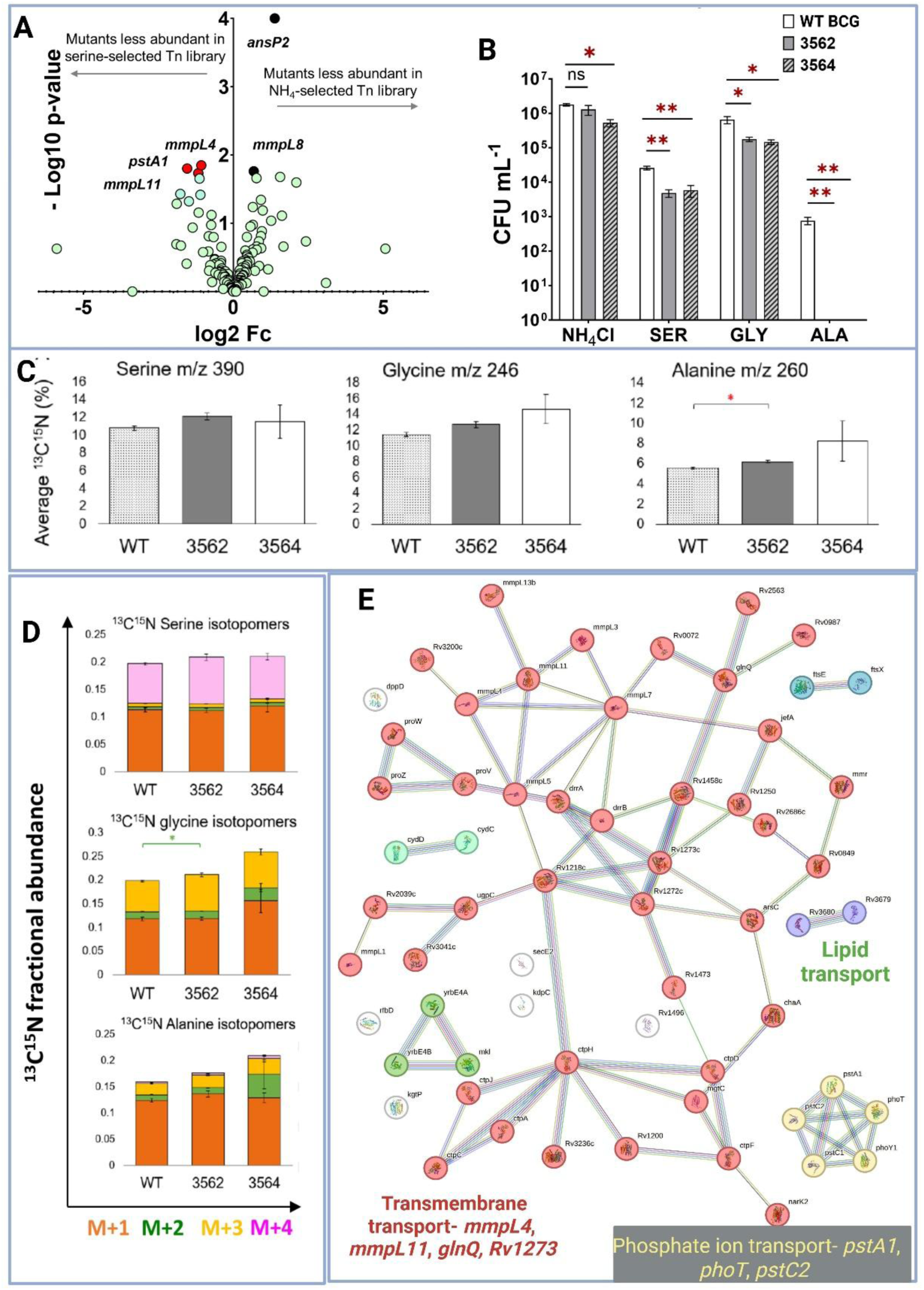
Transposon sequencing identifies candidate Ser transporters in Mtb. **A)** Volcano plot comparing mutant abundance in SER- versus NH₄Cl-selected libraries. The y-axis shows the -log₁₀ *P* value, and the x-axis shows the log₂ fold change (SER vs. NH₄). **B)** Growth of *Mycobacterium bovis* BCG *cycA* Tn mutants on NH₄Cl, serine (SER), glycine (GLY), and alanine (ALA) as sole nitrogen sources, measured as CFU mL^-1^ on Roisin’s minimal medium. Mutants with Tn insertions at positions 3562 and 3564 exhibited significantly reduced growth on SER and GLY and no detectable growth on ALA. * indicate statistically significant differences relative to the WT strain. Data represent the mean of three independent experiments ± SEM, and significance was determined using one-way ANOVA with Dunnett’s multiple-comparison test. **C)** Average ^13^C^15^N % isotopic incorporation in serine, glycine and alanine of WT vs Δ*cycA* mutants. Data are mean ± SEM, with t-test statistical significance (p ≤ 0.05). **D)** Analysis of ^13^C^15^N traced 3-phosphoglyceric acid (PGA) derived amino acids in WT against Δ*cycA* deficient mutants 3562, and 3564. The fractional abundance of mass isotopomers, depending on the number of carbon and nitrogen atoms in the amino acids are shown: serine, 4 isotopomers; glycine, 3 isotopomers; alanine, 4 isotopomers. Mass isotopomers are colour coded. Data are mean ± SEM, with t-test statistical significance (p ≤ 0.05). **E)** STRING v12 protein–protein association network highlighting three major transporter classes lipid transporters, transmembrane transporters, and phosphate ion transporters that showed reduced mutant abundance during growth on SER compared with NH₄Cl, suggesting potential involvement in Ser transport. Line thickness reflects confidence in the predicted associations.

*In silico* annotation identifies *cycA* (Rv1704c) as a possible D-serine/alanine/glycine transporter, yet our Tn-seq data did not support differential selection of Δ*cycA* mutants. To test this directly, we assessed *M. bovis* BCG Δ*cycA* Tn mutants for growth on Ser, Gly, and Ala as nitrogen sources (Fig. 5B). These experiments did not validate *cycA* as essential for Ser or Gly utilisation, as the mutants grew robustly on both nitrogen sources. BCG does not grow reliably on Ala as the sole nitrogen source due to mutations in *ald* gene, which encodes alanine dehydrogenase; therefore, Ala utilisation by the mutants could not be assessed with confidence. Additionally, we performed isotopic labelling experiments using [^13^C^15^N-serine] to assess Ser uptake in the mutants (Fig. 5D). If *cycA* were the primary Ser transporter, we would expect no isotopic incorporation or ^13^C^15^N-MID in the mutants. Instead, the ^13^C^15^N-MIDs were indistinguishable between the WT and the mutants, demonstrating that *cycA* is not a dedicated Ser transporter.

The absence of a clear Ser transporter candidate suggests that either the relevant transporter is essential on 7H11 medium and therefore absent from the library, or that Ser uptake is functionally redundant, with multiple transporters compensating for one another. Consistent with this, very few nutrient transporters were essential in our selections or in the 7H10 dataset of de Jesus et al. (2017)^23^, supporting the idea of transporter redundancy for Ser uptake.

Protein-protein interaction analysis revealed several transporter clusters enriched in the SER-selected library, including transmembrane transporters, ABC-type and ATP-binding transporters, antibiotic-transport proteins, and lipid transporters (Fig. 5E). A subset of transporter mutants was less abundant in the SER-selected library, including the transmembrane proteins *mmpL4* (Rv0450c) and *mmpL11* (Rv0202c), the drug-transport protein Rv1273c, ABC transporters *pctA1* (Rv0931) and *pstC2* (Rv0929), and the glutamine transporter *glnQ* (Rv2564).

In summary, our screen identified several transporters with altered fitness on Ser that warrants further investigation as potential Ser uptake systems and, consequently, possible drug targets. Given that Ser biosynthesis is essential for Mtb’s survival and that host-derived Ser is not accessible to intracellular Mtb^4^, targeting Ser biosynthetic enzymes such as *serC* together with Ser transport pathways may provide effective combinatorial therapeutic strategies. Overall, the Tn-seq screen highlights genes required for Ser metabolism and reveals interconnected vulnerabilities in PYR and folate metabolism that represent promising metabolic drug targets in Mtb.

## Discussion

Ser metabolism has emerged as a central metabolic node in Mtb, with *serC* (Rv0884c) validtaed as a key enzymatic determinant of intracellular survival and virulence^4,15,16^. Previous studies identified *serC* as essential for nitrogen metabolism and *de novo* serine biosynthesis, proposing it as a druggable vulnerability in Mtb^4,15,28^. Our work expands this framework by validating the essentiality of *serC* across multiple infection models and by defining its broader metabolic role in coordinating carbon and nitrogen flux in Mtb.

Across primary human macrophages, RAW 264.7 cells, and *in vivo* infection of BALB/c mice, the Δ*serC* mutant exhibited a profound intracellular growth defect, demonstrating that SerC-dependent serine biosynthesis is indispensable for Mtb replication within host environments. Host metabolism further constrains Mtb’s reliance on this pathway. In mammals, Ser is a non-essential amino acid that supports biosynthesis, one-carbon flux, metabolic homeostasis, and responses to oxidative stress and infection^29^. Dysregulation of Ser biosynthetic enzymes is associated with diverse pathologies, including cancer, diabetes, fatty-liver disease, and neurological disorders. Consistent with this, our previous work showed that Ser availability is significantly reduced in infected macrophages, with more than a two-fold decrease in intracellular Ser pools and diminished flux through the PG3-derived Ser-Gly-Cys pathway^4^. The inability of Δ*serC* Mtb to compensate for this host-driven Ser limitation highlights the pathogen’s vulnerability to Ser scarcity within the intracellular niche.

The complete rescue of the Δ*serC* growth defect by exogenous Ser and Gly highlights the specificity of the *serC*-mediated step, whereas the partial rescue by Thr and Val suggests more indirect metabolic connectivity. These observations align with the central role of the Ser-Gly-one-carbon network in maintaining redox balance, amino-acid homeostasis, and nucleotide synthesis in other bacterial systems^30,31^. Using stable-isotope tracing, we show that loss of *serC* disrupts global carbon and nitrogen metabolic architecture in Mtb. The Δ*serC* strain exhibited reduced incorporation of carbon and nitrogen into multiple biosynthetic pathways, with impaired synthesis of Ser, Gly, Met, Pro, and His amino acids. These findings indicate that *serC* functions not only as a biosynthetic enzyme but also as a metabolic hub supporting broader amino-acid biosynthesis.

Our ^13^C-MFA analysis established that the loss of *serC* reshapes CCM in Mtb. Fluxes toward Ser-derived metabolites including Gly and one-carbon folate-cycle intermediates were elevated in the mutant, consistent with compensatory activation of one-carbon metabolism, to support nucleotide, amino-acid, and phospholipid synthesis. These changes occurred alongside reduced glycolytic, TCA-cycle and MCC fluxes and increased Val degradation and enhanced oxidative PPP flux. The metabolic plasticity of Mtb revealed in our study is consistent with earlier work demonstrating that Mtb possesses a highly flexible metabolic network capable of co-metabolizing cholesterol or glycerol with two-carbon substrates without compartmentalization^4^. The reduced MCC flux observed in the Δ*serC* strain is similar to the low MCC flux profile during growth on cholesterol/acetate, suggesting that Ser biosynthesis might intersect with PROPCoA metabolism and lipid biosynthesis. The increased Val degradation in Δ*serC* suggests the dependence on branched-chain amino acids to supplement PROPCoA pools. Together, these metabolic flux profiles highlight extensive rewiring in the Δ*serC* mutant and establish the centrality of Ser biosynthesis to global metabolic homeostasis in Mtb.

The importance of Ser and Gly metabolism is well established in other bacteria. Yishai et al. (2018) showed that engineered *E. coli* can meet its serine and glycine requirements through a reductive glycine pathway using formate and CO_2_^30^. Similarly, Shi et al. (2022) demonstrated that disruption of Gly synthesis activates compensatory sulfur metabolism and alters antibiotic susceptibility^31^. A recent work has demonstrated species-specific regulation of *serC*. Guo et al. (2025) showed that the nucleoid-associated protein NapR activates *serC* in *M. smegmatis* to promote multicellularity, while repressing *serC* in BCG to modulate growth and Ser sensitivity^32^. Wang et al. (2025) further identified NapR as an activator of *ggr*, tuning the intracellular reactive oxygen species (ROS) levels and coordinating oxidative-stress responses in Mtb^33^. An immunomodulatory role for *serC* has also emerged: Cheng et al. (2024) showed that *serC*-dependent D-Ser production under hypoxia suppresses IFN-γ secretion by CD8⁺ T cells via mTORC1 inhibition^18^. Our findings complement this model by demonstrating that *serC* is simultaneously required for the metabolic robustness necessary for intracellular replication. Collectively, these studies indicate that *serC* supports Mtb’s intracellular survival through sustaining metabolic fitness while modulating host immunity during hypoxic stress.

Our Tn-seq analysis identified the genetic determinants that enable Mtb to use Ser as a nitrogen source, revealing a tightly interconnected metabolic network underpinning Ser-dependent growth. Libraries selected on NH₄⁺ or Ser showed strong agreement with established essentiality datasets, validating the robustness of our selections^16,23,24^. L-Ser deaminase (*sdaA*) and components of the GCS were essential in the Ser-selected library. These findings define the genetic circuitry required for Ser assimilation and highlight the centrality of Ser-linked metabolic networks in Mtb physiology, expanding the repertoire of potential therapeutic targets. We anticipated that Tn-seq would identify Ser transporters required for growth when Ser is the sole nitrogen source, yet no clear candidate was identified. The only amino-acid transporter with differential fitness was *ansP2*, whose mutants were enriched not depleted in the Ser-selected library, inconsistent with a role in Ser uptake. This aligns with prior work showing that *ansP2* primarily mediates asparagine import in Mtb^5^. Although *cycA* is annotated as a potential D-Ser/Ala/Gly transporter^34^, neither Tn-seq nor targeted growth assays supported a requirement for *cycA* in serine or glycine utilization, and isotopic-labelling experiments confirmed intact Ser uptake in Δ*cycA* mutants. Protein-interaction analysis in our study identified several transporter families with altered fitness on Ser including ABC transporters, transmembrane transporters, and lipid transporters that needs further investigation. The absence of a definitive transporter suggests that Ser import may occur through an essential transporter absent from the library or, more plausibly, through a redundant network of transport systems consistent with the broader essentiality landscape in which few nutrient transporters are indispensable under standard conditions. The importance of Ser transport has been investigated in other bacteria. In *E. coli*, *sdaC* encodes Ser transporter and is essential during growth on exogenous Ser^35^. Although Mtb lacks a clear *sdaC* homolog, this study highlights the importance of Ser transport and metabolism across diverse species.

Our findings reveal redundancy in Ser uptake and identify transport-linked genes whose fitness depends on Ser availability. Because intracellular Mtb cannot access host-derived serine and relies on *de novo* synthesis, targeting Ser biosynthetic enzymes such as *serC*, alongside transport pathways, may offer a combinatorial therapeutic strategy^4,36^.

## Conclusion

Our findings validate *serC*-mediated Ser metabolism as a key druggable vulnerability in Mtb. Integrating genetic, metabolomic, isotopic tracing, fluxomics and molecular analyses, we show that disrupting Ser biosynthesis destabilizes central carbon and nitrogen networks and sensitizes the pathogen to targeted inhibition. Targeting Ser biosynthesis alone or in combination with transport and associated genetic elements offers a significant potential for effective combination therapies. More broadly, this work establishes that targeting essential metabolic nodes in a pathogen offers a promising strategy for therapeutic innovations required to combat antimicrobial resistance.

## Materials and Methods

| Strains and cell lines | Reference or source | Identifier or catalog |
| --- | --- | --- |
| *Mycobacterium tuberculosis* H37Rv | American Type Culture Collection (ATCC) | ATCC 27294 |
| *M. tuberculosis* H37Rv ΔserC (Rv0884c knockout) strain | Borah et al., 2019 | - |
| *M. tuberculosis* H37Rv ΔserC::SERC | Borah et al., 2019 | - |
| *M. tuberculosis* H37Rv glnA1 (Rv2220 knockout) strain | This work | - |
| *M. tuberculosis* H37Rv transposon library | This work | - |
| *Mycobacterium bovis* BCG Pasteur | ATCC | ATCC 35748 |
| *M. bovis* cycA transposon mutant | VIB – Vlaams Instituut voor Biotechnologie | (BCG_Tn Insertion-3562 and 3564) |
| RAW 264.7 murine macrophage cell line | ATCC | TIB-71 |

### Reagents

Reagents:
| Chemicals, Peptides, and Recombinant Proteins |  |  |
| --- | --- | --- |
| Rosins minimal media | Borah et al., 2023b | N/A |
| Middlebrook 7H9 | Sigma/Aldrich | M0178-500G |
| Oleic acid-albumin-dextrose-catalase | Becton Dickinson | 212351 |
| Glycerol | Sigma-Aldrich | G7893-1L |
| Tyloxapol | Sigma-Aldrich | T8761-50G |
| Hydrochloric acid | Sigma-Aldrich | 258148-500ML |
| Phosphate buffer saline | Sigma-Aldrich | D8537-500ML |
| Ammonium chloride | Sigma-Aldrich | 299251-20G |
| Tert-butyl dimethyl silyl chloride | Merck Life Science | 00942-10ML |
| $^{13}\text{C}^{15}\text{N}$-serine and $^{15}\text{N}$-NH<sub>4</sub>Cl | | |
| $^{13}\text{C}$glycerol | CK isotopes | |
| Software and Algorithms |  |  |
| Isotopomer Network Compartmental Analysis | Young, 2014 | N/A |
| MATLAB | The MathWorks | N/A |
| Agilent 5975 Chemstation Enhanced Data Analysis | Agilent Technologies | N/A |
| Graph Pad Prism | GraphPad Software | N/A |
| Excel | Microsoft | N/A |

### Mycobacterial cultivation

*Mycobacterium tuberculosis* H37Rv wild type (WT) and mutant strains were cultivated on Middle brook7H11 agar and Middlebrook 7H9 broth with 5% (v/v) oleic acid-albumin-dextrose-catalase enrichment supplement and supplemented with 0.5% glycerol (v/v) as the carbon source. Cultures were grown as static solid agar cultures at 37°C or as liquid cultures at 37°C and agitated at 150 rpm^4^. The auxotrophic strain Δ*serC* constructed in our earlier study was grown in the presence of 50µg.mL^-1^ L-serine and 50µg.mL^-1^hygromycin antibiotic^4^. The complemented strain Δ*serC*::SERC (Borah et al., 2019) was selected with antibiotics hygromycin and kanamycin at 50µg.mL^-1^ and 25µg.mL^-1^ concentrations respectively. *M. bovis* BCG WT and cycA transposon mutants (Tn insertions_3562, and 3564) strains used in this study were obtained from VIB Vlaams Instituut voor Biotechnologie. *M. bovis* strains were cultivated in 7H9 and 7H11 media with OADC as described for H37Rv strains.

### Isotopic labelling of Mtb and *M. bovis* BCG

For isotopic labelling assays, strains were grown in Roisins minimal media with the composition- KH_2_PO_4_, 1 g l^−1^; Na_2_HPO_4_, 2.5 g l^−1^; K_2_SO_4_, 2 g l^−1^; ZnCl_2_, 0.08 mg l^−1^; FeCl3, 0.4 mg l^−1^; CuCl_2_, 0.02 mg l^−1^; MnCl_2_, 0.02 mg l^−1^; Na_2_B_4_O_7_, 0.02 mg l^−1^; NH_4_MoO_4_, 0.02 mg l^−1^; MgCl_2_, 0.0476 g l^−1^; CaCl_2_, 0.055 g l^−1^; Tyloxapol, 01% (v/v); Glycerol, 0.5% (v/v)^37^. All isotopic labelling experiments were performed in three biological replicates. We performed single species isotopic labelling assays of the WT and Δ*serC* strains using two tracers: (a) 30% U-[^13^C_3_] glycerol + unlabelled NH_4_Cl and (b) unlabelled glycerol + U-[^15^N_1_] NH_4_Cl (20%). The labelling experiments with BCG WT and *cycA* mutants were conducted using unlabelled glycerol as the carbon source and 20% [^13^C_3_^15^N_1_]-Serine at 10mM as the sole nitrogen source. Strains were grown up to late exponential growth phases and samples were harvested for metabolomics and mass isotopomer analysis.

### Nitrogen utilisation assays of *Mtb and M. bovis* strains

Mtb and *M. bovis* cultures were grown in 7H9 to a mid-exponential phase. Cultures were harvested and washed thrice in phosphate buffered saline (PBS) by spinning at 4000 rpm for 20 minutes. Washed cells were re-suspended in PBS and serially diluted before plating it out on Roisins minimal agar media with 0.5% of glycerol and various amino acids as sole nitrogen sources at a working concentration of 10 mM: alanine, glycine, histidine, isoleucine, leucine, lysine, methionine, phenylalanine, threonine, tryptophan, valine, asparagine, aspartate, glutamine, glutamate, serine, arginine, cysteine, proline, tyrosine, and ammonium chloride. Colony forming units (CFUs) were counted after two-three weeks post incubation at 37°C. Experiment was conducted using three biological replicates.

### Isolation of primary human macrophages

Leukocytes were isolated from whole blood collected from two independent donors. Briefly, whole blood was diluted with phosphate buffer saline (PBS) up to 50mLs. 25mL blood was added to15mL ficoll (Ficoll® Merck product No. F4375) in falcon tubes and spun 400g for 30 minutes at 20°C (acceleration=1, deceleration=2). The middle ring consisting of peripheral blood mononuclear cells (PBMCs) were collected in a fresh tube and diluted with PBS up to 50mLs. PBMCs were spun at 200g for 20minutes at 20°C (acceleration=2, deceleration=2). Supernatant was discarded and the pellet was resuspended in 50mL PBS and washed by spinning at 200g for 20 minutes at 20°C. after two washes, supernatant was discarded, and cells were resuspended in 25mL RPMI. Cells were counted and seeded in plates and flasks (depending on the assay) for monocytes to stick overnight. Next day, floating cells were removed, and fresh media was added to the monocytes. Monocytes were differentiated to M1-like macrophages for seven days with 50ng/mL of granulocyte-macrophage colony-stimulating factor (GM-CSF) (Merck G5035). Post differentiation, macrophages were washed with warm RPMI and replenished with fresh media for subsequent assays.

### Cell culture

Primary human monocytes and macrophages, THP-1 monocytic cell line and RAW 246.7 cells were cultured using RPMI 1640 media (Merck R8758) supplemented with 10% fetal bovine serum (FBS) (Merck F9665). Cells were grown in 5% CO_2_ incubator set at 37°C and 95% humidity.

### Infection of human and RAW macrophages with Mtb

Infection of cell lines and primary macrophages were done following the methods described in Borah at al. (2019)^4^. Mtb WT, mutant and complemented strains were prepared as described in mycobacterial culture methods. Briefly, bacteria were grown to a late exponential phase in 7H9 broth with appropriate antibiotics and serine as needed for strains and harvested for infection. Bacteria were washed three times in PBS to remove any growth media and antibiotics and resuspended in RPMI cell culture media. THP-1, RAW and primary macrophages were seeded at 1 × 10^5^ cells per well in 12-well plates for infection. Bacteria were added to the cells at a multiplicity of infection (MOI) of five. After three-four hours if infection, infected macrophages were washed with warm PBS supplemented with 0.49 mM Mg2+ and 0.68 mM Ca2+ (PBS+) and fresh RPMI media was added. Infected and uninfected (control) cells were incubated static at 37°C and with 5% CO_2_. Cells were incubated to appropriate time points until their harvest and lysis for CFU and viability assays. At various time points post infection, cells were lysed using 0.1% triton X-100, and cell lysate was serially diluted and plated onto 7H11 plates. WT was plated on 7H11 + OADC, Δ*serC* mutant on 7H1 + OADC + 50µg.mL^-1^ serine + 50µg.mL^-1^ hygromycin and complemented Δ*serC*::SerC mutant on 7H11 + OADC + 50µg.mL^-1^ hygromycin + 25µg.mL^-1^ kanamycin agar plates. Plates were incubated at 37°C and colonies were counted after 2-3 weeks of growth.

### Mouse strains used and ethical approval

Mouse experiments were conducted at two sites: UKHSA and Imperial College (IC). For the study at UKHSA, a total of 45 BALB/c mice, aged six weeks, and free from pathogen-specific infection, were obtained from a UK Home Office accredited supplier (Envigo, UK). Animal studies were conducted at the UK Health Protection Agency (UKHSA, Porton Down, Salisbury, UK). Procedures at IC were conducted under the authority of a UK Home Office approved project licence. Group sizes were determined by statistical power calculations (Minitab, version 16) performed using previous data (SD, approximately 0.56) to detect a difference of 1.0 log_10_ in the median number of colony-forming units (cfu) per millilitre. Animals were divided into three experimental groups of 15 mice and housed in groups of five mice. Cages met with the UK Home Office ‘Code of Practice for the Housing and Care of Animals Bred, Supplied or Used for Scientific Procedures’ (December 2014). The housing environment was maintained within a temperature range of 20-24°C and a relative humidity range of 45 to 65%. Access to food and water was *ad libitum* and environmental enrichment was provided. All experimental work was conducted under the authority of a UK Home Office approved project licence that had been subject to local ethical review at UKHSA by the Animal Welfare and Ethical Review Body (AWERB) as required by the ‘Home Office Animals (Scientific Procedures) Act 1986’.

### Mouse infection experiments

Before the start of the experiment animals were randomly assigned to groups and identified using subcutaneously implanted microchips (Plexx, Netherlands) to enable blinding of the analyses. Mtb H37Rv strains: Mtb H37Rv cultures (wild type (WT), Δ*serC* mutant and complemented strain) were grown in 7H11 and 7H9 broth with 10% OADC (oleic acid, albumin, dextrose, catalase) enrichment (BD life sciences) and 0.05% Tween 80 at 37 °C. Actively growing cultures were used for infection of animals. Challenge for each group of animals was by the aerosol route with Mtb strain H37Rv. Animals were challenged using a contained Henderson apparatus in conjunction with an AeroMP control unit as previously described^38^. Aerosol particles generated were delivered to the animals via both nares using a 3-jet Collison nebuliser. The challenge suspension in the nebuliser was adjusted by dilution in phosphate buffered saline to a concentration of between 5 × 10^6^ to 1 × 10^7^ CFU mL^-1^ to deliver the required estimated, retained, inhaled, target dose of 200 CFUs to the lungs of each animal. The suspension of Mtb in the nebuliser was plated onto Middlebrook 7H11 OADC selective agar to measure the concentration and confirm retrospectively that the expected doses had been delivered. Following infection, animal housing was transferred to Advisory Committee on Dangerous Pathogens (ACDP) containment level 3 and housed within flexible film isolators. Animals were monitored daily, and body weight determined at least weekly throughout the study for 21 days. A range of clinical and behavioural parameters were assessed and where necessary used to determine whether the disease related changes met end-point criteria for euthanasia. Mice remained healthy throughout the period of infection. At days 1, 7 and 21 post-challenge, five animals per timepoint were culled using an overdose of sodium pentabarbital via the intraperitoneal route of delivery. A necropsy was performed immediately after confirmation of death. At necropsy, the lung of each animal were removed and placed into sterile Precellys tubes containing ceramic zirconium oxide beads were homogenised in 1 mL of phosphate buffered saline (PBS). Serial dilutions were plated (0.1 mL per plate, in duplicate) onto Middlebrook 7H11 OADC selective agar. After up to 6 weeks of incubation at 37°C, bacterial colonies were counted and duplicate data averaged to measure CFU mL^-1^of viable Mtb in each tissue sample homogenate. Where no colonies were observed, a minimum detection limit was set by assigning an average count equating to 5 CFU mL^-1^. CFUs were calculated, tests for normality were performed and data were analysed using the Mann whitney test to determine significant differences between the bacterial burden measured between the Δ*serC* mutant and control groups using GraphPad Prism version 10 software (GraphPad Prism software 9.0, Inc., San Diego, CA, United States).

### Gas chromatography mass spectrometry (GC-MS) metabolomics and isotopomer tracing

GC-MS and isotopomer analysis was conducted as previously described^37^. Isotopically labelled pseudo-steady state exponential cultures of Mtb H37Rv and *M. bovis* BCG strains were harvested by centrifugation at 4000 rpm for 20 minutes. Spent media was discarded and the pellet was washed twice in PBS. After the final wash, the pellet was hydrolyzed in 6M hydrochloric acid for 24 h at 100°C and the hydrolysate was dried for mass spectrometry analysis under N_2_ gas. Amino acid hydrolysates were dried and derivatized using pyridine and tert-butyldimethyl silyl chloride (TBDMSCl) (Merck). GC-MS metabolomics was performed to measure amino acid pool sizes and for mass isotopomer analysis. Amino acids were analyzed using a VF-5 ms inert 5% phenyl-methyl column (Agilent Technologies) on GC–MS. Mass spectra were extracted using chemstation GC–MS software. Mass spectrometry datasets were corrected for natural isotope effects using the Isotopomer network compartmental analysis (INCA) platform^37,39^. Average ^13^C or ^13^C^15^N fractional abundances in an amino acid was calculated from the fractional abundance of the mass isotopomers in the amino acid fragments. The corrected mass isotopomer distributions (MIDs) were included into the Mtb model for flux estimation using ^13^C-MFA^21^ (Borah et al., 2021).

### Metabolic modelling and ^13^C-MFA

We used our previously published Mtb H37Rv isotopomer metabolic model^21^. This metabolic network model was supplemented with carbon atom transitions and consisted of a total of 90 reactions and 51 exchange reactions. Metabolic flux estimations were performed using INCA, version 2.2^39^. Fluxes were estimated using a non-linear weighted least square fitting approach to calculate net and exchange flux distributions that were the best fit for generation of the isotopic labelling data and biomass constraints^21,39^. A random initial guess and multistart optimisation approach with 200 restarts was used for flux estimations. Flux profiles with the minimum statistically acceptable SSR were considered the best-fit. The goodness of fit of the flux maps was assessed by comparing the simulated and experimental measurements. The uncertainties in the flux estimations were checked using the Monte Carlo analysis function built into INCA. The upper and lower 95% confidence limits of the best-fit flux distributions were calculated using parameter optimisation option in INCA^39^. Fluxes were significantly different if the 95% confidence limits do not overlap.

### Transposon (Tn) library construction

Temperature labile φMycoMarT7 stocks were amplified using *M. smegmatis* as previously described^40^. Mtb H37Rv was grown at 37°C on 7H11 with 10% OADC, or in 7H9 with 10 % OADC and 0.05 % Tween®80. 100mL cultures of H37Rv were grown for 2-3 weeks. Confluent culture was spun down 4000 rpm for 10 minutes and washed three times with MP buffer (50 mM Tris-HCl, pH 7.5, 150 mM NaCl, 10 mM MgSO_4_, 2 mM CaCl_2_) and resuspended in 9 ml MP buffer at 37 °C. Phage was added at 2 × 10^11^ pfu and incubated for overnight shaking at 30 rpm at 37 °C. Next day cultures were washed once with PBS with 0.05 % Tween80 and then resuspended in PBS Tween80. To determine the number of transductants the cultures were serially diluted and plated on 7H11 agar plates with 20 µg/ml kanamycin and 0.05 % Tween80 for 2-3 weeks. The efficiency of transductions was 400,000 CFUs. Colonies were scraped of plates and collected in 7H9 containing 15 % glycerol 0.05 % Tween80 and stored in -80 °C. For selection, tubes were taken out of the storage and thawed and 100µL were diluted and plated on to Roisin’s minimal media with 20 ug/ml kanamycin, 0.05 % Tween®80, 0.5 % glycerol and either 0.5 % NH_4_Cl, or 10 mM serine. The library was enumerated by serial diluting and plating on Roisin’s with both or both 0.5 % NH_4_Cl, and 10 mM serine.

DNA was isolated using phenol:chloroform isolation^40^. The resulting DNA was sheared and blunt ended, added A tail and ligated with adapters. Transposon junctions were amplified using the primers listed in Table S1. To minimise PCR bias during the preparative amplification stage, the number of PCR cycle was optimised. For these reactions a 10 µl PCR was set up that consisted of 5 µl of Q5 High-Fidelity 2× Master Mix (NEB, UK), 0.5 µl × 20 EvaGreen® Dye (Biotium), 0.5 µl IS6 and 0.5 µl of Himar-1 MycoMarT7 mariner transposon (2 pmol µl^-1^ primers), 1 µl of template DNA and 2.5 µl of nuclease free water. The following PCR conditions were used to optimise the preparative amplification: denaturation at 95 °C for 2 min followed by 40 cycles at 95 °C for 10 sec, annealing at 58 °C for 10 sec and extension at 72 °C for 30 sec. The product sizes were analysed using Agilent 2100 Bioanalyser. The optimum number of cycles was the lowest number of cycles that gave maximal amplification. For each sample 4 × 50 µl preparative PCR amplifications were completed using the optimised cycle number. Control reactions with IS6 primer/template only and IS6/MarX primer mix with no template were used. Following the amplification step, samples were pooled and purified using 1× volume AMPure XP® beads and eluted in 50 µl water. The fragment size was assessed using gel electrophoresis on a 0.8 % agarose gel and Agilent 2100 Bioanalyser to check for a desired fragment length of ∼500 bp. DNA concentration was quantified using a Quantus™ Fluorometer (Promega). The purified and indexed samples were mixed in equimolar amounts and sequenced on an Illumina NovaSeq X.

---

**Table S1.**
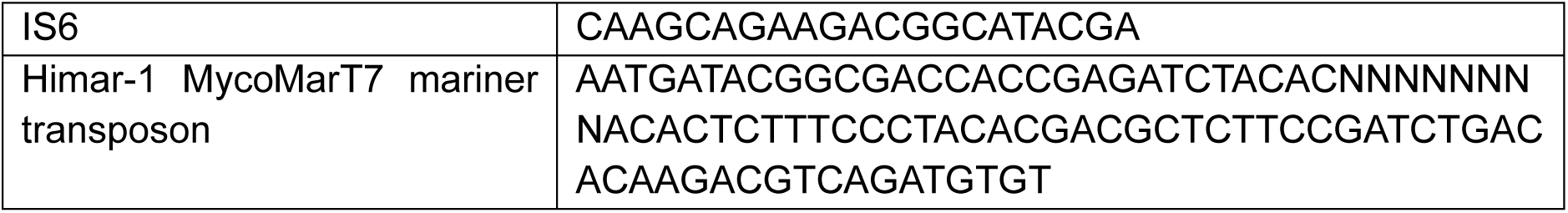
Primers for amplification of transposon junctions.

## Acknowledgements

We are grateful to VIB - Vlaams Instituut voor Biotechnologie for providing us *M. bovis* BCG gltBD transposon mutant (BCG_3922c TnInsertion-8654).

## Author contributions

MJP: conducted investigations, formal analysis, data visualization, writing review and editing

TAM: experimental design, writing review and editing

DK, JS: conducted investigations, writing review and editing

BR, RW, SC: experimental design and animal work, writing review and editing

JM: conceptualized the project, funding acquisition, writing review and editing

KBS: conducted investigations, experimental design, formal analysis, data curation and visualisation, conceptualized the project, funding acquisition, procured resources, writing original draft.

